# Chiral twisting in cytoskeletal polymers regulates filament size and orientation

**DOI:** 10.1101/459974

**Authors:** Handuo Shi, David Quint, Ajay Gopinathan, Kerwyn Casey Huang

**Affiliations:** Department of Bioengineering, Stanford University, Stanford, CA 94305, USA; Department of Physics, University of California at Merced, Merced, CA 95343, USA; NSF-CREST : Center for Cellular and Biomolecular Machines, University of California at Merced, Merced, CA 95343, USA; Department of Microbiology and Immunology, Stanford University, Stanford, CA 94305, USA; Chan Zuckerberg Biohub, San Francisco, CA 94158

## Abstract

While cytoskeletal proteins in the actin family are structurally similar, as filaments they act as critical components of diverse cellular processes across all kingdoms of life. In many rod-shaped bacteria, the actin homolog MreB directs cell-wall insertion and maintains cell shape, but it remains unclear how structural changes to MreB affect its physiological function. To bridge this gap, we performed molecular dynamics simulations for *Caulobacter crescentus* MreB and then utilized a coarse-grained biophysical model to successfully predict MreB filament properties *in vivo.* We discovered that MreB double protofilaments exhibit left-handed twisting that is dependent on the bound nucleotide and membrane binding; the degree of twisting determines the limit length and orientation of MreB filaments *in vivo.* Membrane binding of MreB also induces a stable membrane curvature that is physiologically relevant. Together, our data empower the prediction of cytoskeletal filament size from molecular dynamics simulations, providing a paradigm for connecting protein filament structure and mechanics to cellular functions.

## Introduction

The actin and tubulin families of cytoskeletal proteins constitute essential components of cellular physiology in virtually all bacteria, archaea, and eukaryotes. Despite structural similarities within each of the two families, their primary functions span a diverse range of processes including cell morphogenesis^1^, division^2,3^, and DNA segregation^4^. In bacteria, many of these cytoskeletal proteins form filaments that are highly dynamic *in vivo.* Structural tools such as X-ray crystallography and cryo-electron microscopy have elucidated various filament structures within the bacterial actin family, including anti-parallel, straight double protofilaments of MreB^5^, single, polar polymers of FtsA^3^, and bipolar, anti-parallel filaments of ParM^4^, suggesting that filament conformations are highly tunable and have been selected for particular physiological functions over evolutionary time. However, the links between the conformational dynamics of these proteins *in vivo* and the molecular mechanisms by which they regulate cell physiology remain undiscovered. Molecular dynamics (MD) simulations are a powerful tool for identifying protein structural dynamics and filament mechanics at atomic resolution, providing key information to map filament properties from the protein to the cellular scales.

One such cellular-scale property defined by a bacterial actin homolog is cell shape, which is ultimately dictated by the rigid cell wall, a highly crosslinked mesh of peptidoglycan. During growth, cells actively remodel their cell wall while robustly maintaining their shape^6^. In rod-shaped bacteria such as *Escherichia coli,* cell-wall synthesis during elongation is regulated by the widely conserved actin homolog MreB^7^, which dictates the pattern of insertion of new cell-wall material^8^ and thereby maintains rod shape^7,9^. Genetic depletion and chemical inhibition of MreB lead to misshapen cells and eventually cell lysis^10,11^. Many point mutations in MreB alter cell shape in subtle ways, such as changing cell width^12-14^, curvature^15^, or polar morphology^14,16^ without affecting viability. In *E. coli,* MreB forms short filaments that move along the cell periphery^1^, and the localization and movement of these filaments are correlated with cell width^17^. MreB movement is chiral, which induces twist in the cell body during elongation^17,18^. Previous MD studies of *Thermotoga maritima* MreB (TmMreB) showed that ATP hydrolysis and polymerization affect MreB monomer conformation, which in turn regulates the bending of MreB dimers^19^. The bending of a TmMreB dimer was also altered *in silico* by binding the membrane protein RodZ, which directly interacts with MreB and tunes cell shape^20^. In *E. coli,* MreB forms antiparallel double protofilaments^5^ that can deform membranes^21^, and the double protofilament conformation is essential for rod-shape maintenance in *E. coli^5^.* However, it remains obscure how molecular-level changes in MreB connect to the biophysics of the double protofilament structure, and to the functions of MreB *in vivo.*

In this study, we exploited the recent solution of a crystal structure of a double protofilament of *Caulobacter crescentus* MreB^5^ (CcMreB) to uncover the connection between MreB structural dynamics *in silico* and filament conformation *in vivo.* We performed all-atom MD simulations for each step during CcMreB filament assembly (Fig. 1), from monomers to single protofilaments, and then to double protofilaments with or without a membrane. Simulations of double protofilaments revealed a new left-handed twisting conformation in ATP-bound double protofilaments. The degree of twisting was reduced when the double protofilaments were bound to ADP or a membrane, and binding to a membrane induced membrane curvature mimicking that of bacterial cells. We used our MD simulations to extract parameters relevant for coarse-grained analyses of membrane-bound MreB double protofilaments, from which we established a connection between intrinsic twisting and filament limit length, which we verified *in vivo* with *E. coli* MreB mutants. Taken together, our results link the molecular-scale behaviors of MreB to cellular phenotypes in *E. coli,* providing a paradigm for connecting protein structure to cellular function across disparate length scales.

**Figure 1:**
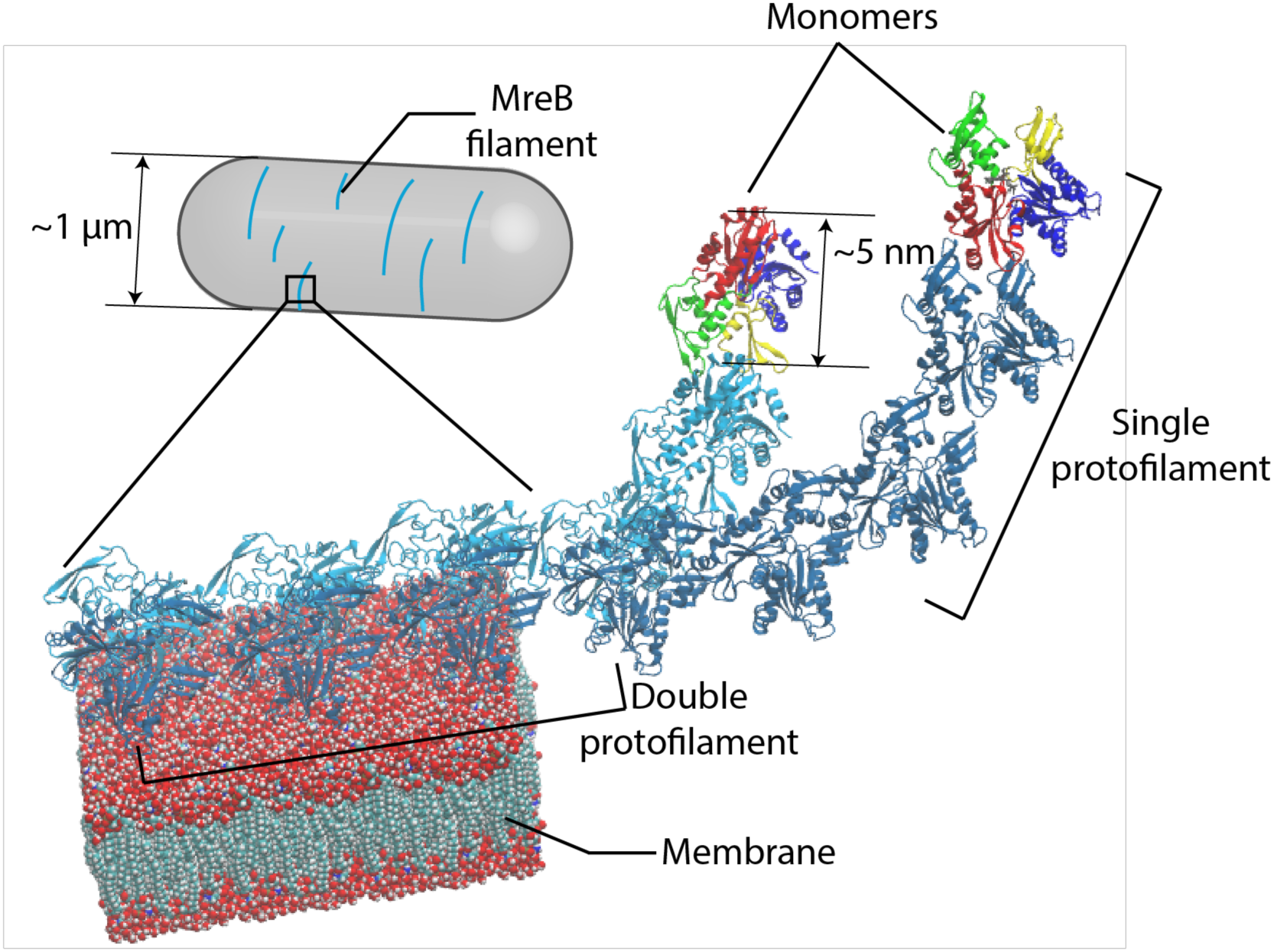
Assembly of MreB protofilaments. MreB monomers first polymerize into single protofilaments. Next, two antiparallel single protofilaments assemble into a double protofilament, with membrane-binding domains on the same side of the double protofilament^5^. Inside bacterial cells, short MreB filaments bind the inner face of the plasma membrane, align approximately circumferentially, and rotate around the long cell axis to guide cell-wall insertion and to determine rod-like shape and size.

## Results

### MreB monomer conformation is nucleotide- and polymerization-dependent

To study the first step of MreB oligomeric assembly (Fig. 1), we performed all-atom MD simulations of MreB as a monomer and as a dimer in a single protofilament (Fig. 2a, Methods). All simulations were initialized from the crystal structure of the CcMreB single protofilament (PDB ID: 4CZF)^5^. By analogy with actin, we refer to the two subunits in an MreB dimer as the (+) and (−) ends (Fig. 2a, right). The four subdomains were defined by aligning the MreB structure to that of actin, with the nucleotide bound in the center of the four subdomains (Fig. 2b)^5^.

**Figure 2:**
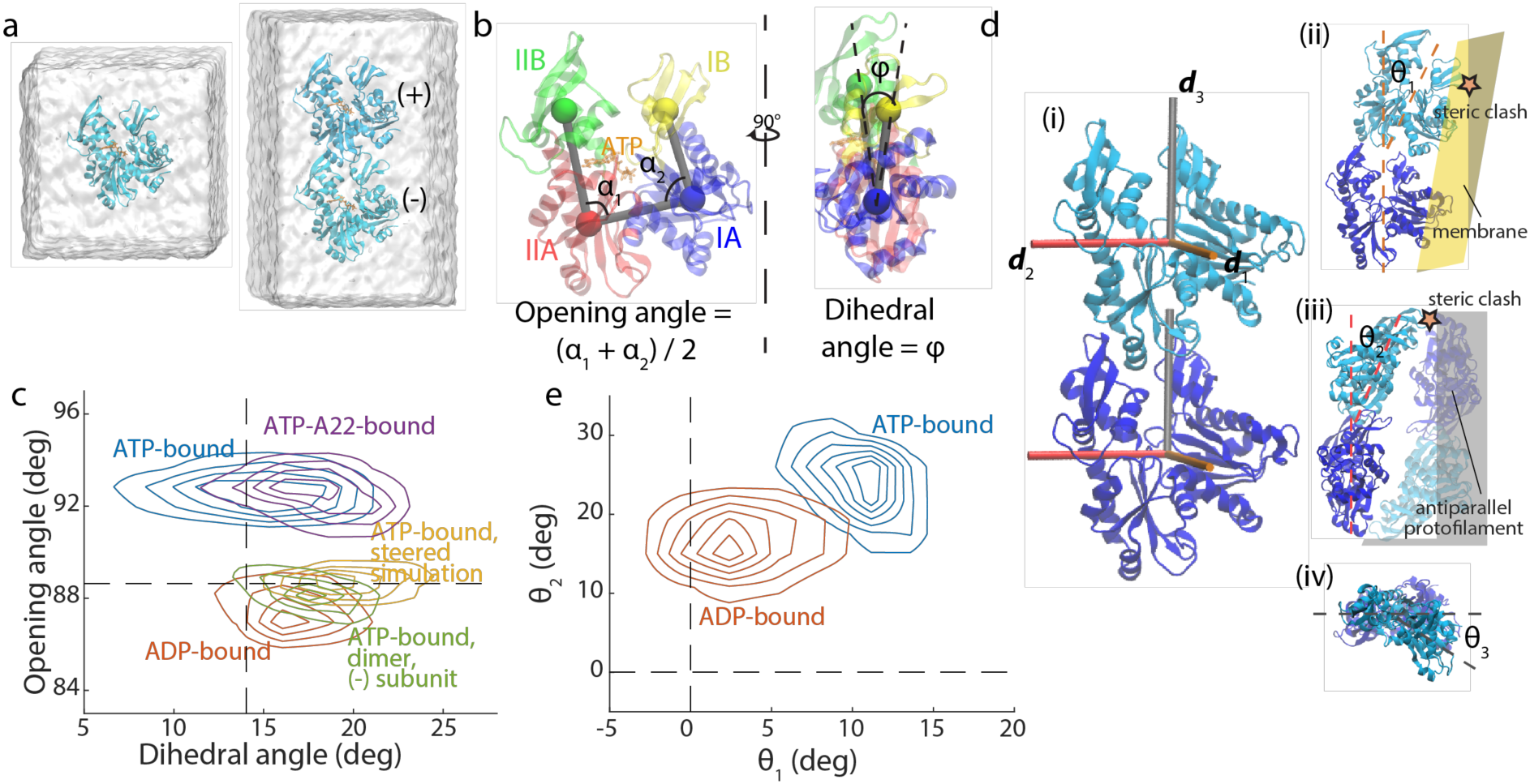
MreB monomer and dimer conformations are nucleotide-dependent. a. Simulated systems of an MreB monomer (left) and a single protofilament with two subunits (“2×1 protofilament”, right). Each MreB subunit is bound to a nucleotide, with the whole system surrounded by a water box. In the 2×1 single protofilament, we refer to the top and bottom MreB subunits as the (+) and (−) subunits, respectively.
b. Definitions of opening angle and dihedral angle for an MreB monomer, with the centers-of-mass of the four subdomains shown as colored spheres.
c. Contour density plot of the distributions of opening and dihedral angles for each simulation system from the last 40 ns of the simulation. MreB subunits essentially adopted one of two conformations in simulations. ATP-bound MreB monomers exhibited large opening angles in the presence (purple) and absence (blue) of A22, while an ADP-bound monomer (red) and the (-) subunit of an ATP-bound dimer (green) had smaller opening angles. Steering of the opening angle of an ATP-bound monomer to its value in the crystal structure (yellow) mimicked the conformation of an ADP-bound monomer. Dashed lines denote the values of the opening and dihedral angles in the crystal structure (PDB ID: 4CZF).
d. (i) An MreB dimer from a single protofilament, with three axes overlaid on each subunit that were used to compute the degree of bending and twisting between them. (ii) Illustration of 01, with positive 01 denoting bending toward the membrane surface (yellow). Positive 01 leads to a steric clash with the membrane surface. (iii) Illustration of 02, with positive 02 denoting bending toward the interprotofilament interface. The paired antiparallel protofilament is shown in semitransparency. Positive 02 leads to a steric clash with the paired protofilament. (iv) Illustration of 03 from the top of a protofilament, with positive 03 denoting lefthanded twisting.
e. Contour density plot for the distributions of 01 and 02 from the last 40 ns (200 frames) of the simulations, with both ATP- and ADP-bound single protofilaments bending toward the membrane side and toward the inter-protofilament interface. An ATP-bound single protofilament exhibited more substantial bending in both directions than an ADP-bound single protofilament. Dashed lines denote the respective angles in the crystal structure.

In our simulations of an ATP-bound MreB monomer, we observed a rapid opening of subdomains IB and IIB, exposing the ATP-binding pocket. We quantified conformational changes by measuring the angles formed by the centers-of-mass of the four subdomains, defining an in-plane opening angle and an out-of-plane dihedral angle (Fig. 2b). The ATP-bound MreB monomer adopted a more open state, with an opening angle of ∼92° at the end of an 80-ns simulation, compared to the ∼88° opening angle of an ADP-bound MreB monomer (Fig. 2c, Fig. S1a). The dihedral angle was slightly smaller in an ATP-bound monomer than in an ADP-bound monomer (Fig. 2c, Fig. S1 b), consistent with the larger dihedral angle in the crystal structure of CcMreB bound to ADP (PDB ID: 4CZL) versus CcMreB bound to AMP-PNP (PDB ID: 4CZM) (Fig. S1c). This result qualitatively differed from our previously reported MD simulations using TmMreB, in which ATP-bound TmMreB exhibited larger dihedral angle than ADP-bound TmMreB but a similar opening angle^19^. To interrogate this difference, we performed new simulations using ATP-bound TmMreB, and obtained results consistent with our previous study (Fig. S1d,e)^19^. Therefore, although CcMreB and TmMreB are structurally similar, they likely adopt different conformations upon nucleotide binding. Such observations may relate to polymeric differences observed *in vitro,* wherein TmMreB formed straighter protofilaments on rigid lipid tubes than CcMreB^5^.

We next asked how MreB conformation *in silico* is affected by the MreB inhibitor S-(3, 4-dichlorobenzyl) isothiourea (A22) by performing MD simulations with MreB bound simultaneously to both ATP and A22 (Methods). Although A22 is known to perturb cell morphology *in vivo* by targeting the active site of MreB^15,22^, the molecular mechanism of action is still obscure. In our simulations, A22 did not affect the MreB monomer opening angle, and only slightly increased the dihedral angle (Fig. 2c). Thus, our results suggest that A22 does not directly affect MreB monomer conformation and is unlikely to alter the ATP-binding pocket, consistent with other studies proposing that A22 blocks phosphate release rather than inhibiting ATP hydrolysis^5,23^.

By contrast to the open conformation of an ATP-bound monomer, the (-) subunit in an ATP-bound MreB dimer maintained a closed conformation resembling the ADP-bound monomer (Fig. 2c), a conformation similar to subunits within a CcMreB protofilament crystal structure (Fig. S1c)^5^. We calculated opening and dihedral angles for all published CcMreB crystal structures^5^, and found that monomeric structures have larger opening angles than polymerized structures (Fig. S1c), supporting our conclusion that polymerization closes MreB.

Motivated by previous findings relating MreB conformation to ATP-binding pocket stability^19^, we quantified ATP-binding pocket stability by calculating the buried solvent-accessible surface area (SASA) between MreB and ATP (Methods). Buried SASA quantifies the surface area of an ATP-MreB interface (Fig. S1f), and thus a larger buried SASA indicates a more stable ATP-binding pocket. The buried SASA in an ATP-bound CcMreB monomer decreased coincident with increases in opening angle (Fig. S1g,h), and ATP-A22-bound CcMreB and ATP-bound TmMreB exhibited similar decreases (Fig. S1g). By contrast, the (-) subunit of an ATP-bound CcMreB dimer maintained high buried SASA, indicating that its ATP-binding pocket remained stable. To verify that the buried SASA of ATP is related to the opening angle, we performed steered simulations of an ATP-bound CcMreB monomer in which we constrained the opening angle to the crystal structure value of ∼89°. Although the dihedral angle opened slightly in the steered simulation (Fig. S1a,b), the buried SASA of ATP remained high (Fig. S1g). Similarly, in our TmMreB monomer simulations, we observed a similar reduction in buried SASA when TmMreB opened (Fig. S1i). Taken together, our simulations suggest that CcMreB monomers adopt distinct open and closed conformations; ATP-bound CcMreB monomers prefer the open state but close upon polymerization. The closed state may facilitate ATP hydrolysis by increasing the stability of the ATP-binding pocket.

### Bending of an MreB single protofilament is nucleotide-dependent

We next sought to study the conformational changes in single protofilaments with two CcMreB subunits (“2×1 protofilaments”) by analyzing the relative movements of the (+) and (-) subunits in the dimer (Fig. 2a,d). We simulated CcMreB 2×1 protofilaments with both subunits bound to ATP or ADP, and quantified their relative orientation changes by calculating the Euler angles that characterize the three orthogonal modes of rotation around the *x, y,* and *z* axes (Fig. 2d(i)): *θ*_1_ and *θ*_2_ characterize bending into the membrane surface and inter-protofilament surface, respectively (Fig. 2d(ii, iii)), and *θ*_3_ characterizes twisting along the protofilament (Fig. 2d(iv)). We defined all three Euler angles to be zero in the crystal structure (Fig. 2d(i)). A stable membrane-binding double-protofilament conformation requires *θ*_1_ to be negative and *θ*_2_ to be approximately zero to avoid steric clashes (Fig. 2d(ii,iii)). We found that the largest changes in our simulations occurred in the bending angles (Fig. 2e, Fig. S1j,k), whereas no systematic protofilament twisting was observed (Fig. S1l). The bending angles were also nucleotide-dependent, with ATP-bound protofilaments exhibiting larger bending angles than ADP-bound protofilaments (Fig. 2e), consistent with our previously reported results in TmMreB^19^.

Considering the double protofilament structure and the membrane binding interface (Fig. 2d(ii,iii)), both bending angles observed in our 2×1 protofilament simulations are unlikely to occur in a double-protofilament architecture. A non-zero θ_2_ would destabilize the inter-filament interface (Fig. 2d(iii)) and split the double protofilament. Positive θ_1_ corresponds to bending toward the membrane surface (Fig. 2d(ii)), whereas *in vitro* experiments indicate that MreB filaments bend away from the membrane^5^. Therefore, although single-protofilament simulations demonstrate the molecule’s capacity for nucleotide-dependent conformations, simulations of a double protofilament conformation and consideration of membrane binding are critical for revealing MreB structural dynamics that are relevant *in vivo.*

### ATP-bound MreB double protofilaments twist in a membrane-dependent fashion

We next performed MD simulations of MreB double protofilaments, each containing four MreB doublets (a 4×2 protofilament, Fig. 3a). Simulations were performed with all MreB subunits bound to ATP or ADP (Fig. 3a,b, Methods), and at least two replicate simulations were performed for all systems. Similar to our analysis of 2×1 protofilaments (Fig. 2), we quantified the three Euler angles for neighboring doublet pairs in the double protofilaments. To minimize boundary effects, we first focused on the middle doublet (pair 2; Fig. 3a). As expected, bending of each protofilament was dramatically different in a double protofilament versus a single protofilament. *θ*_1_ values were smaller in magnitude and were generally negative (Fig. S2a), indicating slight bending away from the membrane, and *θ*_2_ decreased to approximately zero (Fig. S2b). Instead of bending along *θ*_2_, which would disrupt the symmetry and stability of a double protofilament, twisting (*θ*_3_) was prominent in the double protofilament (Fig. 3c, Fig. S2c). In all 4×2 protofilament simulations, left-handed twisting was observed. Interestingly, in water, an ATP-bound double protofilament twisted more (10.3±2.1°, mean±S.D. from Gaussian fitting of last 40 ns of simulation) than an ADP-bound double protofilament (4.2±2.0°), suggesting that the difference in *θ*_2_ bending between ATP- and ADP-bound single protofilaments was resolved into double protofilament twisting. To confirm that our observations on bending and twisting were not artefacts due to limited filament size, we performed a larger simulation with eight ATP-bound MreB doublets in water (an 8×2 protofilament). In this 60-ns simulation, changes in bending and twisting angles matched our observations in 4×2 protofilaments (Fig. S2d-f, Movie S1). To verify that the double-protofilament twist was not unique to CcMreB, we constructed a homology model of *E. coli* MreB (Methods), and found that EcMreB exhibited quantitatively similar left-handed twisting in simulation (Fig. S2g). Thus, higher-order oligomerization can dramatically alter the biophysical properties of MreB filaments.

**Figure 3:**
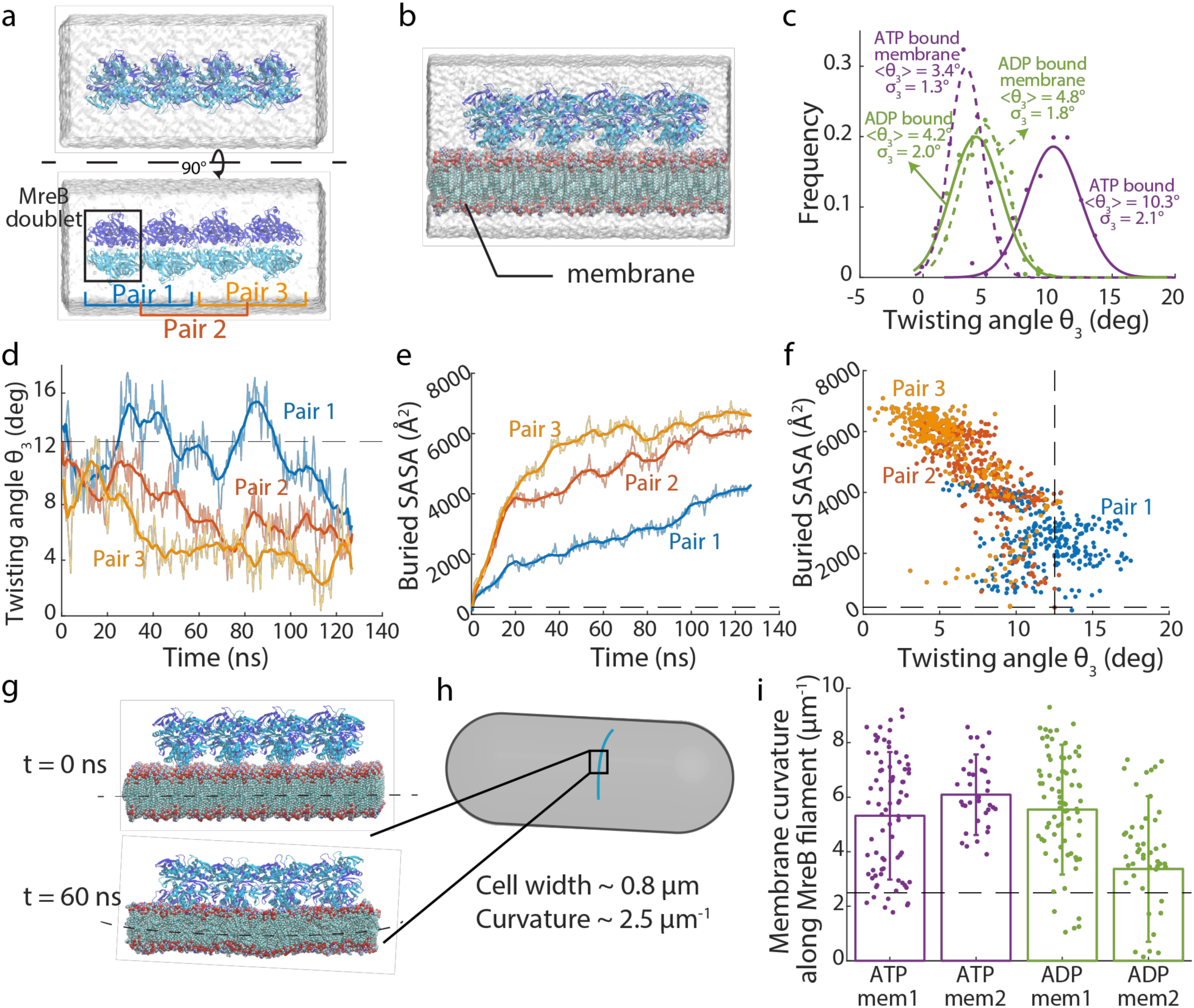
Binding of an MreB double protofilament to the membrane decreases MreB twist and induces membrane curvature. a. Simulated system of a 4×2 MreB double protofilament in water. The system consists of four MreB doublets (eight subunits), surrounded by a water box.
b. Simulated system of a 4×2 MreB double protofilament bound to a membrane. The MreB protofilament was placed near a membrane patch, with the membrane-binding side of the double protofilament (with subdomains IA and IB) facing the membrane patch. The spaces on the top and bottom of the membrane patch not occupied by MreB were filled with water.
c. Distribution of twisting angles in simulated systems at equilibrium. At the start of the simulations, all systems had zero twisting. The ATP-bound 4×2 protofilament displayed a large twisting angle, which was reduced when the 4×2 protofilament bound the membrane. Membrane binding did not substantially affect the twisting angle of ADP-bound protofilaments. Solid dots are histograms from the last 40 ns (200 frames) of each simulation, and curves are Gaussian fits of the histograms. The mean and standard deviation for each Gaussian fit are also indicated.
d. Twisting angles lessened over time when a pre-twisted 4×2 protofilament was placed close to a membrane patch. Untwisting occurred first in Pair 3, then propagated to Pair 2 and Pair 1. The dashed line shows the initial twisting angle in Pair 2.
e. Buried SASA of the membrane-binding interface for the twisted protofilament. Higher buried SASA indicates stronger membrane interaction. Similar to the changes in twisting angles, the buried SASA increased first in Pair 3, then in Pair 2 and Pair 1. The dashed line is the initial buried SASA for Pair 2.
f. Scatter plot of buried SASAs and twisting angles in the simulation analyzed in (d,e). Each dot represents the values for a certain Pair at a particular time point, and dashed lines are the initial values for Pair 2. Buried SASA and twisting angle were highly correlated (Pearson’s *r* = 0.79, *p* < 10^-10^, Student’s f-test).
g. In a typical membrane simulation with 4×2 protofilaments, the membrane started flat (top) and ended up curved toward MreB (bottom).
h. MreB filament orientation inside a bacterial cell. MreB binds the inner face of the cytoplasmic membrane, with the membrane curving toward MreB filaments.
i. Values of induced membrane curvature in 4×2 protofilament simulations are comparable with *in vivo* membrane curvatures. The dashed line represents the *invivo* reference value for a rod-shaped cell with width 0.8 μm. Data points represent the mean ± standard deviation for the last 40 ns of each simulation.

A twisted double protofilament is not compatible with binding to a flat membrane. To address this incompatibility, we performed MD simulations of 4×2 protofilaments in the presence of a membrane patch (Fig. 3b). Membrane binding reduced twist in ATP-bound double protofilaments but did not affect the less-twisted ADP-bound structures (Fig. 3c). To test the hypothesis that membrane binding suppresses twisting in ATP-bound double protofilaments, we took the twisted protofilament structure from the end of an ATP-bound 4×2 protofilament simulation in water, and placed it ∼10 Å away from a membrane patch. Within 120 ns, the filament untwisted from one end to the other (Fig. 3d, Movie S2), effectively “zippering” into the membrane. The decrease in twist angle from each doublet was accompanied by an increase of buried SASA in the protein-membrane interface, indicative of stronger MreB-membrane interactions (Fig. 3e,f). Therefore, membrane binding directly suppresses twisting in ATP-bound MreB double protofilaments.

We further asked whether membrane binding alters the stability of the double protofilament conformation, as quantified by the distances between the interacting V118 residues within each MreB doublet (Fig. S2h), which are essential for forming a double-protofilament structure^5^. For both ATP- and ADP-bound double protofilaments, our simulations in water exhibited increased distances between V118 residues in the first 60 ns (Fig. S2i), suggesting a destabilized double protofilament interface. In contrast, membrane-associated simulations maintained short V118 distances (Fig. S2i), indicating more stable double protofilaments. Therefore, membrane binding potentially stabilizes the double-protofilament structure.

### Double protofilaments induce physiologically relevant membrane curvatures

The distinct structures of MreB double protofilaments when bound or unbound to a membrane patch and the lack of complete untwisting when membrane-bound (Fig. 3c) indicated that membrane binding introduced strain into the MreB filaments that may affect membrane conformation. In our simulations, the membrane started flat, but after ns, the membrane bent toward the MreB protofilaments (Fig. 3g). In rod-shaped bacterial cells, the membrane also bends toward MreB filaments, forming a curvature dictated by the cell width (Fig. 4h). We computed the curvature at the center of the membrane patch along the protofilament direction and found that the membrane curvatures for all 4×2 protofilament membrane simulations were ∼5 μm^-1^ (Fig. 3i), on the same scale as the membrane curvature of a rod-shaped bacterial cell that is ∼0.8 μm in width (∼2.5 μm^-1^).

**Figure 4:**
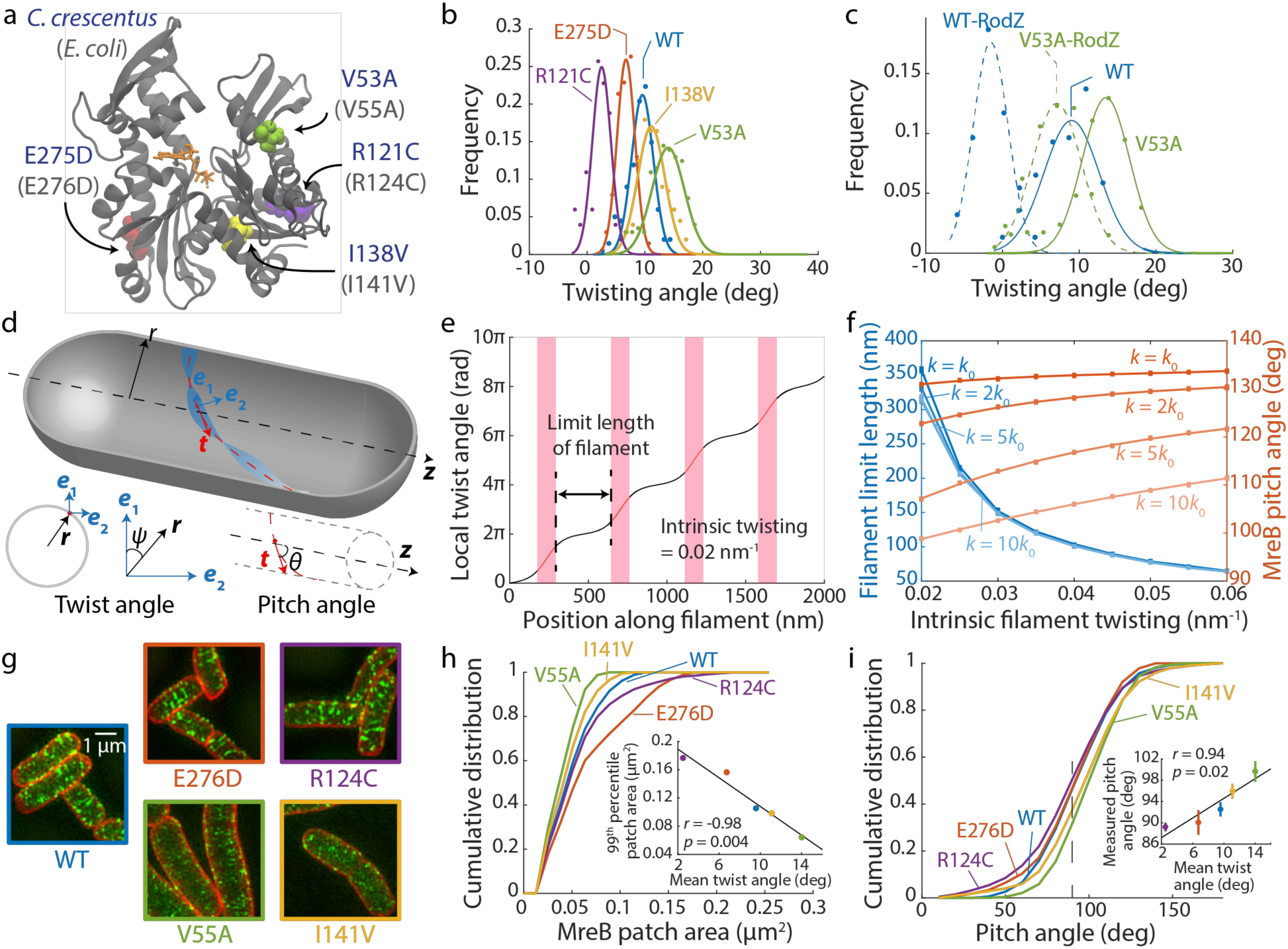
MreB twisting angle predicts MreB filament limit length *in vivo.* a. Mutations in MreB investigated via MD simulations mapped onto the CcMreB crystal structure. These mutations were previously identified to alter cell shape^14,20^, and are conserved between CcMreB (residue numbers in blue) and EcMreB (residue numbers in light gray). Colored spheres: mutated residues. Orange: ATP molecule. Gray: MreB protein structure.
b. Distributions of twisting angles in simulations of CcMreB mutants. All systems started with zero twisting. E275D and R121C twisted less than wild-type MreB, while V53A and I138V twisted more. Dots are histograms from the last 40 ns of each simulation, and curves are Gaussian fits of the histograms.
c. Distributions of twisting angles with and without RodZ binding. For wildtype and the V53A mutant, binding of the cytoplasmic tail of RodZ decreased twisting. The effect of RodZ binding was approximately additive to the effects of MreB mutation, such that the V53A-RodZ system twisted more than the WT-RodZ system. Dots are histograms from the last 40 ns of each simulation, and curves are Gaussian fits of the histograms.
d. Schematic of our coarse-grained model. The MreB filament (blue) is bound to the inside of a cylindrical cell body (gray) with radius *r.* The centerline of the filament is drawn in red, with ***t*** being the tangent vector. ***e***_1_ and ***e***_2_ are the material frame coordinates along the filament. Local twisting angle *ψ* is defined as the angle between ***e***_1_ and the unit vector ***r***. Local pitch angle *θ* is defined as the angle between tangent vector ***t*** and the cylindrical centerline.
e. The coarse-grained model predicts that when a filament with a given intrinsic twist bound to a cylindrical membrane, the filament forms flat domains (black lines) interspaced with twisted 360° turns (red lines in pink shading). It is energetically favored for the filament to break at the twisted regions and thereby form only flat fragments, so we predict that the extent of a flat domain sets the limit length of a membrane-bound filament.
f. The coarse-grained model predicts that filaments with larger intrinsic twisting have shorter limit length. Similarly, the coarse-grained model predicts that the orientation of a short filament (90% of the corresponding limit length) deviates more from 90° as the intrinsic twist increases. Increasing the intrinsic bending *k* did not affect the limit length, but reduced the pitch angle to be closer to 90°. Data points are mean ± standard error of the mean from 20 independent Monte Carlo simulations, and the smoothed curves are fit to a third-order polynomial as a guide to the eye. For most data, the error bars are small and overlap with the data points.
g. Structured illumination microscopy of wildtype and the four EcMreB mutants constructed in *E. coli* cells with a sandwich fusion of msfGFP to MreB. Images are maximum projections of a z-stack, with red (membrane dye FM 4-64FX) and green (MreB-msfGFP) channels merged.
h. The cumulative distributions of MreB-msfGFP fluorescence patch sizes for each strain trend with the twisting angles in (b). The V55A and I141V strains had smaller patch sizes than wildtype, and E276D and R124C strains contained larger patches. MreB patches were defined as continuous regions with high msfGFP signal on the cylindrical cell body with area larger than the diffraction limit. *n* > 1,000 patches were measured for each strain. Inset: the 99^th^ percentile of patch area in each strain was highly correlated with the mean twist angle from (b) (Pearson’s *r* = −0.98, *p* = 0.004, Student’s t-test, *n* = 5 strains), providing experimental validation of the coarse-grained model.
i. Cumulative distributions of MreB filament pitch angle. The V55A and I141V strains had larger pitch angles than wildtype, while E276D and R124C had smaller pitch angles that were closer to 90°. The pitch angle was defined as the angle between the main axis of each fluorescent patch and the long axis of the cell. Inset: the experimentally measured pitch angle highly correlated with the mean twist angle from (b) (Pearson’s *r* = 0.94, *p* = 0.02, Student’s t-test, *n* = 5 strains). Data points are mean ± standard error of the mean for *n >* 1,000 patches in each strain.

To validate that the observed membrane curvature changes were related to the twisted nature of 4×2 protofilaments, we performed simulations of 2×1 protofilaments in the presence of a membrane patch as a control. The membrane patches bound to 2×1 protofilaments were more variable and did not exhibit a characteristic curvature throughout the simulation (Fig. S2j). Thus, only double MreB protofilaments induce stable and physiologically relevant curvature in the membrane, suggesting that MreB needs to form double protofilaments for its function *in vivo.*

### Mutation of MreB and binding of the regulatory protein RodZ modulate intrinsic twist

We hypothesized that since many MreB mutations alter cell shape, they potentially also induce altered intrinsic twist and membrane interactions as a double protofilament. We identified four MreB mutants that were reported to cause a range of alterations to *E. coli* cell shape, with the corresponding residues conserved between CcMreB and EcMreB: R124C^24^, E276D^20^, A55V^14^, and I141V^14^. The four mutated residues are spread across the MreB structure (Fig. 4a), and thus potentially alter MreB function in different manners.

We first performed all-atom MD simulations for each of the corresponding CcMreB mutants bound to ATP in a 4×2 protofilament configuration in water. All mutants exhibited similar bending (Fig. S3a,b), but differed widely in twisting angles compared to wild-type CcMreB: E275D (E276D in EcMreB) and R121C (R124C in EcMreB) twisted less than wildtype, whereas V53A (V55A in EcMreB) and I138V (I141V in EcMreB) exhibited more twist (Fig. 4b, Fig. S3c).

We then asked whether these mutants also exhibit differential twisting when membrane-bound by simulating 4×2 protofilaments of R121C and V53A in proximity to a membrane patch. These two mutants were selected because they exhibited the smallest and the largest intrinsic twisting in our MD simulations in water, respectively (Fig. 4b). Despite the large differences in intrinsic twisting of these mutants in water, they behaved similarly when bound to a membrane, where twist angles were suppressed down to similar levels as wild-type MreB (Fig. S3d-f). Therefore, genetic perturbations can modulate the intrinsic twist of MreB double protofilaments without disrupting the ability of MreB to form stable membrane-binding complexes or to maintain rod-shaped growth. However, to untwist a highly twisted filament costs more energy compared to a less twisted filament, which potentially alters the conformation or orientation of membrane-bound MreB *in vivo.*

The membrane protein RodZ directly interacts with MreB^25^ and is essential for rod-shape maintenance^26^. *E. coli* cells actively tune the stoichiometry of MreB and RodZ as a function of growth rate and growth phase^20,27^, and changes in the MreB:RodZ ratio alter the localization pattern of MreB and cellular dimensions^20^. We previously showed that RodZ binding and MreB mutations that complement the loss of rod-like shape in *ΔrodZ* cells both alter the mechanics of single TmMreB protofilaments *in vivo*^20^. Therefore, we hypothesized that RodZ binding also affects MreB double-protofilament conformations. We constructed a homology model for the cytoplasmic tail of *C.crescentus* RodZ from the co-crystal structure of *T. maritima* RodZ and MreB (PDB ID: 2UWS)^25^, and aligned it to the RodZ-binding interface for each of the subunits in a 4×2 CcMreB protofilament (Methods). We then performed all-atom MD simulations of the system in water, and found that while RodZ binding did not substantially change either of the bending angles in a double protofilament (Fig. S3g,h), it significantly reduced the twisting angle of MreB (Fig. 4c, Fig. S3i). As the ratio of MreB and RodZ in *E. coli* cells varies from ∼10:1 to ∼4:1 depending on growth conditions^20^, our simulations suggest that RodZ abundance actively regulates MreB filament conformation *in vivo*^20^.

Since MreB mutations and RodZ binding both alter the twisting of a MreB double protofilament, we further performed MD simulations for an MreB mutant (V53A) bound to the cytoplasmic tail of RodZ; the V53A 4×2 protofilament in the absence of RodZ exhibited the largest twisting in our simulations (Fig. 4b). Simulations of RodZ bound to a V53A 4×2 protofilament exhibited partially suppressed twisting (Fig. 4c, Fig. S3i), with an average twist slightly lower than that of wild-type MreB in the absence of RodZ (Fig. 4c). The additivity of effects on twisting suggests that RodZ and MreB mutations can alter double protofilament twist in orthogonal manners. Therefore, although regulatory proteins are likely to modulate the intrinsic twisting in MreB double protofilaments, they likely shift the absolute twist but keep the order of twist angles across mutants.

### MreB twisting angle predicts filament-limit length and pitch angle *in vivo*

How does the intrinsic twist of MreB double protofilaments affect MreB conformation *in vivo*? To answer this question, we utilized a coarse-grained model^28^ in which an MreB double protofilament is represented as a beam, with its bending and twisting stiffness extracted from our all-atom MD simulations (Methods). Considering that the large turgor pressure across the bacterial cell envelope (∼1 atm^29^) forces the membrane to adopt a shape matching that of the cell wall, we treated the membrane as a rigid cylindrical surface. We calculated the Hamiltonian for an infinitely long MreB beam with intrinsic twist and bend^28^, and identified the local twist and bend angles that minimize its energy (Methods, Fig. 4d). Intuitively, in the presence of a binding interaction between the filament and the membrane, a twisted filament can gain binding energy by untwisting so that more of its membrane-binding interface can bind the membrane, but the untwisting process also accumulates bending and twisting energy. Therefore, competition between membrane binding and filament mechanics ultimately determines the minimal-energy conformation, which involves periodic flat (untwisted) domains along the filament that are bound to the membrane^28^. For an infinitely long filament, these flat regions are separated by short regions of unbinding that introduce a local twist of 2π (Fig. 4e), relieving the accumulated twist energy. However, in a protein filament with a finite subunit-subunit interaction energy, it could be energetically more favorable to introduce a break in the filament rather than retain a twist wall between successive flat regions that cannot bind to the membrane. The energetic cost for breaking an MreB filament (i.e. eliminating two intrafilament monomer bonds) can be roughly estimated as the energy of hydrolyzing two ATP molecules (∼40 *k_B_T*). This cost can easily be compensated for by the ensuing membrane binding of the twist regions, as the twist regions are generally tens of nm long (Fig. 4e) and contain ∼40 MreB monomers, each with an affinity of ∼10 *k_B_T*^30^. Thus, since it is energetically favorable for the twist walls to be absent, leaving only finite flat regions bound to the membrane, we predicted that MreB filament lengths *in vivo* are limited to be shorter than each flat domain.

The coarse-grained model predicts that the limit length of MreB filaments should decrease with increasing intrinsic twisting (Fig. 4f). Similarly, the local pitch angle θ (Fig. 4d) balances between filament bending and twisting: with a pitch angle of 90°, the filament fully untwists but largely preserves bending; when the pitch angle deviates from 90°, the filament reduces bending while remaining somewhat twisted. Therefore, from an energetic point of view, our coarse-grained model predicts that the intrinsic twisting in an MreB filament (which we define to be 90% of the limit length) causes its orientation to deviate from the perfect circumferential direction (pitch angle *θ* = 90°) (Fig. 4f). We further performed sensitivity analyses by altering the parameters that affect filament conformation^28^. For instance, by varying the intrinsic bending *k*, we find that the limit-length predictions are largely unaffected, whereas larger values of *k* lead to pitch angles closer to 90° (Fig. 4f). Similarly, altering the ratio of bending and twisting moduli (*C/K*) changes the pitch angle but not limit length (Fig. S3j), while decreasing membrane binding potential decreases the limit length without affecting the pitch angle (Fig. S3k). Notably, despite variation in the predicted values across parameters, our model generally predicts that larger intrinsic twist leads to short filaments with larger pitch angles.

To verify the results of our coarse-grained model, we experimentally constructed *E. coli* strains expressing the MreB mutants (Fig. 4a) with a sandwich fusion of monomeric super-folder green fluorescent protein (msfGFP)^31^ as the sole copy of MreB. To quantify the shape and size of the MreB filaments, we imaged each strain using super-resolution structured illumination microscopy (Methods). In wild-type cells, MreB formed short filaments with a limit length of ∼200-300 nm (Fig. 4g), approximately consistent with the prediction of our coarse-grained model (Fig. 4f). The E276D and R124C mutants clearly contained much longer filaments that spanned roughly half the cell periphery, whereas V55A and I141V had very short MreB filaments (Fig. 4g). We quantified the distribution of MreB patch areas in each mutant as a proxy for filament length, and indeed E276D and R124C had larger MreB patches than wildtype, and V55A and I141V had smaller patches (Fig. 4h). We used the 99^th^ percentile of patch size as an approximation for filament limit length in each mutant, and found that it was highly negatively correlated with the twisting angles we observed in all-atom MD simulations (Fig. 4h, inset), consistent with our coarse-grained model. Similarly, we calculated the pitch angle of each MreB patch from the microscopy images (Fig. 4i) and observed that MreB filament orientation positively correlated with intrinsic twist (Fig. 4i, inset): a larger intrinsic twist led to a larger deviation from circumferential orientation. Taken together, our microscopy results validated the predictions of our coarse-grained model that the intrinsic twist of MreB double protofilaments affects filament limit length and orientation *in vivo.*

## Discussion

Here, we used MD simulations to reveal a new twisted double-protofilament conformation of CcMreB (Fig. 3c) and EcMreB (Fig. S2g). We determined that twisting is regulated by various factors including the binding nucleotide (Fig. 3c), the membrane (Fig. 3c), genetic perturbations (Fig. 4b), and regulatory proteins (Fig. 4c). While previous MD simulations of TmMreB provided insights into the structural properties of MreB at the monomer and single-protofilament levels^19,20^, the twist only occurs with a double-protofilament structure. Using a coarse-grained model, we further linked the intrinsic twisting of MreB filaments to their size limit and orientation when bound to the membrane (Fig. 4e,f). Since EcMreB shares a higher sequence similarity with CcMreB (62%) than with TmMreB (52%), our MD studies in CcMreB also permit more versatile mutagenesis studies linking simulations to experimental measurements in *E. coli,* from which we validated our coarse-grained model *in vivo* with fluorescence measurements of MreB mutants predicted to have altered twist (Fig. 4g-i).

Twisting of MreB breaks symmetry and introduces chirality. Chirality is a common feature of biological systems: chiral asymmetry during embryogenesis ensures the normal function of the heart, gut, and brain^33^, the spirals of snail shells generally exhibit right-handed chirality^34^, and the tendrils in climbing plants also grow with specific chirality^35^. In bacterial growth, chirality has been observed at the population^36^ and single-cell^17,18^ levels, and can be altered by perturbing MreB or other components of the cell-wall synthesis machinery^17^. Our simulations have for the first time revealed a molecular-level mechanism for the origin of chirality (Fig. 3c), with handedness that is consistent with that of single-cell twisting in *E. coli*^17,18^. Further understanding of the emergence of asymmetry and MreB twisting will benefit from recent advances in protein design^37^. The design of MreB mutants with various intrinsic twists can be directly tested *in vivo* to further probe the connections between molecular twisting and single-cell physiology. The observation that RodZ alters MreB twist (Fig. 4c) suggests that a host of other proteins that may similarly tune MreB conformation, whose expression may variably impact cell shape under various growth conditions. Further, general rules dictating filament twisting can be utilized to construct synthetic architectures in cells that have variable binding interfaces, mechanical properties, and, as we have shown for MreB, tunable lengths and orientations when bound to a membrane.

Much remains to be learned about the links among MreB, its regulatory partners, and cell-wall synthesis. Our prediction that binding of MreB double protofilaments induces physiologically relevant membrane curvature (Fig. 3g-i) is at least qualitatively consistent with electron microscopy of purified MreB bound to *in vitro* membranes^21^, and may be important for geometric localization of MreB^8^. The induced membrane curvatures are slightly larger than the curvature of bacterial cells, potentially due to the limited size of our simulation system and the lack of turgor pressure in our simulation. While *in vitro* assays of MreB’s interaction with the membrane are challenging due to its N-terminal amphiphilic helix, further coarse-grained approaches incorporating the mechanical properties of the membrane and turgor pressure will further broaden our understanding of MreB’s role in geometric sensing and cell-shape determination. While previous models have studied how MreB orientation is related to filament mechanics^30,38^, they have either assumed a non-twisted filament conformation^30^, or neglected the fact that membrane binding only occurs on a specific side of the filament^38^. Therefore, our coarse-grained model provides a more comprehensive view of MreB mechanics and ultrastructure.

Beyond MreB, many other bacterial actin homologs such as FtsA, ParM, and MamK also polymerize into filaments. While these proteins have diverse roles in bacteria, our study suggests that nucleotide binding and protein-protein interactions may generally induce conformational changes in these polymers whose discovery can be accelerated with MD simulations. Despite their common structural homology to actin, these proteins exhibit diverse protofilament architectures^39^, which may reflect their varied physiological roles from cell division to plasmid segregation. That binding of RodZ or genetic mutations in MreB altered or even reversed chirality (Fig. 4e,h) reflects remarkable flexibility in the intrafilament interface of MreB, wherein single mutations can exert enormous impact on mesoscopic filament conformation and cell shape. Chirality reversal in mammalian cells distinguishes cancerous cells from normal cells, and such chirality is dependent on the functionality of the actin cytoskeleton^40^. Moreover, modulation of chirality is not limited to the actin family: single mutations can also introduce twist to filaments of the bacterial tubulin homolog FtsZ, resulting in growth along a helical pattern rather than a ring^41^. Thus, understanding the molecular origin of chirality in cytoskeletal filaments has broad implications for studying chiral morphogenesis and identifying potential factors that alter or reverse chirality.

## Methods

### Equilibrium MD simulations

All simulations were performed using the MD package NAMD^42^ with the CHARMM36 force field^43^, including CMAP corrections^44^. Water molecules were described with the TIP3P model^45^. Long-range electrostatic forces were evaluated by means of the particle-mesh Ewald summation approach with a grid spacing of <1 Å. An integration time step of 2 fs was used^46^. Bonded terms and short-range, non-bonded terms were evaluated every time step, and long-range electrostatics were evaluated every other time step. Constant temperature (*T* = 310 K) was maintained using Langevin dynamics^47^, with a damping coefficient of 1.0 ps^-1^. A constant pressure of 1 atm was enforced using the Langevin piston algorithm^48^ with a decay period of 200 fs and a time constant of 50 fs. Setup, analysis, and rendering of the simulation systems were performed with the software VMD^49^. Steering of the opening angle was achieved by introducing collective forces to constrain the angle to defined values through the collective variable functionality of NAMD^42^.

### Simulated systems

MD simulations performed in this study are described in Table S1. Unless otherwise noted, systems were initialized from the crystallographic structure of *C. crescentus* MreB bound to magnesium and ADP (PDB ID: 4CZF)^5^. The bound nucleotide was replaced by ATP or ADP with chelating Mg^2+^ ions for all simulated systems. In simulations including a membrane, patches consisting of phosphatidylethanolamine (POPE) were generated using the membrane plugin in VMD. Water and neutralizing ions were added around each simulated system, resulting in final simulation sizes of up to 480,000 atoms. For mean values and distributions of measurements, only the last 40 ns were used for each simulation. All simulations were run until equilibrium was reached unless specified in the text. To ensure simulations had reached equilibrium, measurement distributions were fit to a Gaussian.

### Analysis of dihedral and opening angles

The centers-of-mass of the four subdomains of each protein subunit were obtained using VMD, excluding the amphiphilic helix (residues 1 to 8). For each time step, we calculated one opening angle from the dot product between the vector defined by the centers-of-mass of subdomains IIA and IIB and the vector defined by the centers-of-mass of subdomains IIA and IA. Similarly, we calculated a second opening angle from the dot products between the vectors defined by the centers-of-mass of subdomains IA and IB and of subdomains IA and IIA. The opening angles we report are the average of these two opening angles (Fig. 2b, left). The dihedral angle was defined as the angle between the vector normal to a plane defined by subdomains IA, IB, and IIA and the vector normal to a plane defined by subdomains IIB, IIA, and IA (Fig. 2b, right).

### Calculation of bending and twisting angles in single and double protofilaments

At each time step of a simulation, the coordinate system of the bottom and top subunits (or each subunit pair) was defined using three unit vectors (***d_1_***, ***d_2_***, ***d_3_***)^50^. For single protofilaments, d3 approximately aligns to the center of mass between the two subunits, ***d_2_*** is defined to be perpendicular to the membrane plane, and ***d_1_*** = ***d_3_*** × ***d_2_*** (Fig. 2d). The same definitions for the unit vectors were used for double protofilaments. The rotation angle around ***d_3_*** (*θ*_3_) represents twist between the bottom and top subunits (or subunit pair). Similarly, rotations around ***d_2_*** and ***d_1_*** (*θ*_2_ and *θ*_1_) represent bending parallel to the membrane plane and bending toward the membrane plane, respectively (Fig. 2d).

### A22 force field generation

The A22 structure was isolated from PDB ID 4CZG using UCSF chimera^51^ by removing all other molecules and adding missing hydrogens in the original PDB file. The force field file for A22 was generated using SwissParam with default parameters^52^.

### Calculation of buried solvent-accessible surface area (SASA)

The interaction strength between two interacting molecules was estimated by calculating the contact surface area between them, which can be approximated by measuring the surface area buried between the two molecules that is not accessible to solvent when the molecules interact. This surface area is known as the buried SASA. The buried SASA between two molecules can be calculated from three quantities: the SASA of each molecule by itself (denoted as *A*_1_ and *A*_2_), and the SASA of the complex of the two molecules when interacting (denoted as *A*_1+2_). If the molecules are in contact, then the sum of the SASA of each molecule is greater than the SASA for both molecules together, and the contact area is the difference between the two values divided by two (to account for double counting):

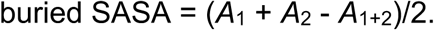

### Construction of homology models for *E. coli* MreB and *C. crescentus* RodZ

Homology models were constructed using the software MODELLER^53^. Using MreB as an example, the amino acid sequences of EcMreB and CcMreB were aligned using the UniProt website (http://www.uniprot.org/aliqn/). The alignment results and the PDB file with the CcMreB crystal structure were processed by MODELLER to generate 10 homology models. The homology model with the lowest DOPEHR score was used for MD simulations.

### Calculation of membrane patch curvature in simulations

The positions of each phosphate atom in the top layer of the membrane (the layer that directly interacts with MreB) were extracted and fit to a second-order polynomial. The curvature of the membrane patch was defined as the curvature at the center of the fitted surface.

### Coarse-grained simulations

The Hamiltonian of the filament is^28^

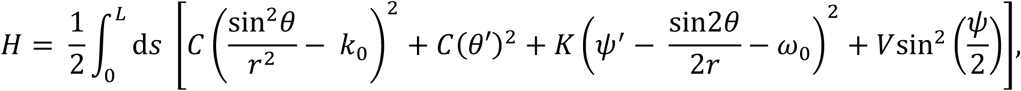

where *L* is the total length of the filament, *θ* and *ψ* are the local tilt and twist angles, respectively, *r* is radius of the cell, *C* is the bending modulus of the filament, *K* is the torsional modulus, *V* is the membrane binding potential, and *k*_0_ and ω_0_ are the intrinsic bending and twisting of the filament, respectively. Parameter values are listed in Table S2.

The total energy per unit length was minimized for an infinite-length filament bound to an infinitely long cylinder by searching for solutions that are periodic over an arc distance *l*. The boundary conditions were set to be

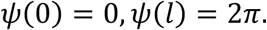

The Hamiltonian was then minimized with respect to *θ*, *ψ*, and *l*, yielding both the equilibrium period *l* and the equilibrium filament shape described by *θ* and *ψ*.

The energy was computed by discretizing the Hamiltonian into *N* segments, with each segment *i* able to adopt a distinct bending and twisting conformation described by angles *θ_i_* and *ψ_i_*. The discretized Hamiltonian was used to calculate the total energy of the filament as

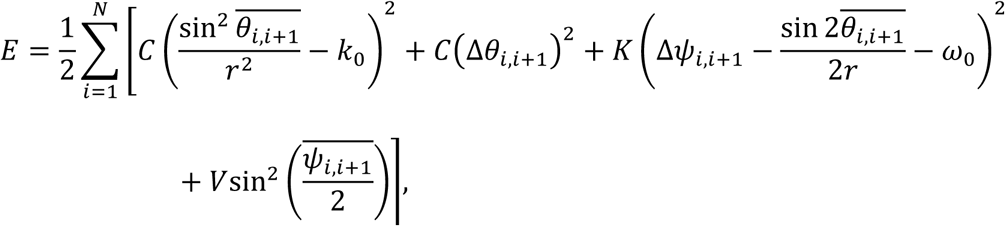

where 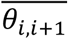 and 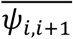 are the average tilt and twist angles between nearest neighbor segments, and Δ*θ*_*i,i*+1_ and Δψ_*i,i*+1_ are the differences in tilt and twist angles between nearest neighbor segments. A classical Metropolis Monte Carlo algorithm was used to minimize the energy of the system. Specifically, for each *l*, starting from an initial configuration of *θ* = 90° and *ψ’* = 2π/*l*, each Monte Carlo step *t* altered *θ_i_* or *ψ_i_* to change the filament conformation from *z^t^* to a trial conformation *z*’. The new conformation *z*^(*t*+1)^ was determined using the Metropolis algorithm:

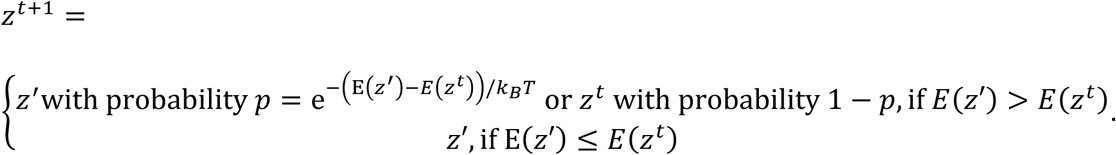

Results were assessed to have converged after ∼10^7^ Monte Carlo steps, as defined by energy fluctuations lower than 1% of the minimized energy across the last 10^4^ steps. The corresponding period *l* leading to the minimized energy was identified using a Golden-section search. Twenty independent replicate simulations were carried out for each parameter set to ensure that a global minimum was reached.

### Estimation of parameters for coarse-grained modeling

The bending and torsional moduli of MreB filaments were estimated from the variance of the appropriate simulations. For the torsional modulus *K*, the standard deviation, σ, of the fluctuations in the twist angle from 4×2 protofilament simulations was ∼1.88° per monomer length. From this value, the torsional rigidity can be estimated as *K* = *k_B_TΔ/σ*^2^, where *Δl* ∼ 5 nm is the length of an MreB monomer. This estimate gives *K* ∼ 4.6×10^3^ *k_B_T* nm. The bending modulus *C* can be estimated similarly. Genetic mutations in MreB did not substantially alter *K* or *C*. The membrane-binding potential of each MreB monomer was estimated to be 10 *k_B_T* in a previous study^30^, yielding *V* = 4 *k_B_T*/nm for a double protofilament. See Table S2 for a list of all parameters used in our coarse-grained simulations.

### Strains and media

Strains used in this study are listed in Table S3. All strains were grown with aeration at 37 °C in LB medium (10 g/L tryptone, 5 g/L yeast extract, and 5 g/L NaCl).

### Sample preparation and imaging for structured illumination microscopy

Saturated overnight cultures were back-diluted 1:200 into pre-warmed fresh LB and grown at 37 °C with shaking. The cultures were further diluted 1:10 into pre-warmed fresh LB at 60 min and 150 min after the first dilution, respectively. By 220 min, the cultures reached exponential growth with OD∼0.1. One milliliter of the cells was fixed in phosphate-buffered saline containing 3% glutaraldehyde/3% paraformaldehyde (Electron Microscopy Sciences) at room temperature for 15 min, with 1 μg/mL FM 4-64FX membrane stain (Invitrogen) added during fixation. Cells were washed three times in cold phosphate-buffered saline, and 1 μL of the cell solution was pipetted onto a No. 1.5 coverslip (Zeiss) coated with poly-L-lysine solution (Sigma-Aldrich). After the droplet dried, a small drop of ProLong Diamond AntiFade Mountant (Thermo Fisher) was added on top of the dried droplet, and the coverslip was mounted on a glass slide (VWR) and sealed with VALAP (equal parts Vaseline, lanolin, and paraffin by weight).

Cell samples were imaged on an OMX V4 microscope platform (GE Life Sciences) with a 100X (NA 1.42) oil-immersion objective (Nikon Instruments). Images from two channels were collected on two Evolve 512 electron-multiplying charged couple device cameras (Photometrics) using DeltaVision microscopy imaging system v. 3.70 (GE Life Sciences).

### Image analysis for structured illumination microscopy

Raw images were reconstructed and aligned into 3D z-stacks using SoftWoRx v. 6.5.2 (GE Life Sciences). The middle plane for each z-stack was segmented by the FM 4-64FX signal using *Morphometries*^54^ to obtain individual cell contours. For each contour, a coordinate-system mesh was calculated using the pill mesh function from *MicrobeTracker^55^.* A three-dimensional surface was reconstructed from the segmentation mesh assuming rotational symmetry about the central axis, and MreB patches localized near the cell periphery were identified from the GFP channel based on intensity, with patches smaller than the diffraction limit for structured illumination microscopy (∼0.02 μm^2^) excluded from quantification.

## Supplementary Information

The supplementary information contains 3 figures, 3 tables, and 2 movies.

## Acknowledgments

The authors thank the Huang lab and Zemer Gitai for helpful discussions. This work was supported in part by an Agilent Graduate Fellowship and a Stanford Interdisciplinary Graduate Fellowship (to H.S.), National Science Foundation (NSF) grant DMS-1616926 (to A.G.), NSF CAREER Award MCB-1149328, NIH Director’s New Innovator Award DP2-OD006466, and the Allen Discovery Center at Stanford University on Systems Modeling of Infection (to K.C.H.). K.C.H. is a Chan Zuckerberg Biohub Investigator. AG and DQ were supported by the NSF-CREST: Center for Cellular and Bio-molecular Machines at UC Merced (NSF-HRD-1547848). The authors also acknowledge the hospitality of the Aspen Center for Physics, which is supported by NSF grant PHY-1607611. Structured illumination microscopy in this study was supported, in part, by Award Number 1S10OD01227601 from the National Center for Research Resources. The contents of this study are solely the responsibility of the authors and do not necessarily represent the official views of the National Center for Research Resources or the National Institutes of Health.

## Author Contributions

H.S. and K.C.H. conceptualized the study. H.S., D.Q., A.G., and K.C.H. designed the experiments. H.S. and D.Q. performed simulations. H.S. created strains, carried out imaging experiments, and analyzed the data. H.S. and K.C.H. wrote the manuscript. All authors reviewed the manuscript prior to submission.

## Data Availability

The datasets generated during the current study are available from the corresponding author on reasonable request.

## Author Information

The authors have no competing financial interests. Correspondence and requests for materials should be addressed to.

## References

1 Van Teeffelen, S. et al. The bacterial actin MreB rotates, and rotation depends on cell-wall assembly. Proceedings of the National Academy of Sciences 108, 15822-15827 (2011).

2 Adams, D. W. & Errington, J. Bacterial cell division: assembly, maintenance and disassembly of the Z ring. Nature Reviews Microbiology 7, 642-653 (2009).

3 Szwedziak, P., Wang, Q., Freund, S. M. & Löwe, J. FtsA forms actin-like protofilaments. The EMBO journal 31, 2249-2260 (2012).

4 Gayathri, P. et al. A bipolar spindle of antiparallel ParM filaments drives bacterial plasmid segregation. Science 338, 1334-1337 (2012).

5 Van den Ent, F., Izoré, T., Bharat, T. A., Johnson, C. M. & Löwe, J. Bacterial actin MreB forms antiparallel double filaments. Elife 3, e02634 (2014).

6 Holtje, J. V. Growth of the stress-bearing and shape-maintaining murein sacculus of Escherichia coli. Microbiol Mol Biol Rev 62, 181-203 (1998).

7 van den Ent, F., Amos, L. A. & Löwe, J. Prokaryotic origin of the actin cytoskeleton. Nature 413, 39-44 (2001).

8 Ursell, T. S. et al. Rod-like bacterial shape is maintained by feedback between cell curvature and cytoskeletal localization. Proceedings of the National Academy of Sciences 111, E1025-E1034 (2014).

9 Jones, L. J., Carballido-Löpez, R. & Errington, J. Control of cell shape in bacteria: helical, actin-like filaments in Bacillus subtilis. Cell 104, 913-922 (2001).

10 Wachi, M. et al. Mutant isolation and molecular cloning of mre genes, which determine cell shape, sensitivity to mecillinam, and amount of penicillin-binding proteins in Escherichia coli. Journal of Bacteriology 169, 4935-4940 (1987).

11 Bean, G. et al. A22 disrupts the bacterial actin cytoskeleton by directly binding and inducing a low-affinity state in MreB. Biochemistry 48, 4852-4857 (2009).

12 Harris, L. K., Dye, N. A. & Theriot, J. A. A. Caulobacter MreB mutant with irregular cell shape exhibits compensatory widening to maintain a preferred surface area to volume ratio. Molecular microbiology 94, 988-1005 (2014).

13 Monds, R. D. et al. Systematic perturbation of cytoskeletal function reveals a linear scaling relationship between cell geometry and fitness. Cell reports 9, 1528-1537 (2014).

14 Shi, H. et al. Deep phenotypic mapping of bacterial cytoskeletal mutants reveals physiological robustness to cell size. Current Biology 27, 3419-3429. e3414 (2017).

15 Dye, N. A., Pincus, Z., Fisher, I. C., Shapiro, L. & Theriot, J. A. Mutations in the nucleotide binding pocket of MreB can alter cell curvature and polar morphology in Caulobacter. Molecular microbiology 81, 368-394 (2011).

16 Kawazura, T. et al. Exclusion of assembled MreB by anionic phospholipids at cell poles confers cell polarity for bidirectional growth. Molecular microbiology 104, 472-486 (2017).

17 Tropini, C. et al. Principles of bacterial cell-size determination revealed by cell-wall synthesis perturbations. Cell reports 9, 1520-1527 (2014).

18 Wang, S., Furchtgott, L., Huang, K. C. & Shaevitz, J. W. Helical insertion of peptidoglycan produces chiral ordering of the bacterial cell wall. Proc Natl Acad Sci U S A 109, E595-604, doi:10.1073/pnas.1117132109 (2012).

19 Colavin, A., Hsin, J. & Huang, K. C. Effects of polymerization and nucleotide identity on the conformational dynamics of the bacterial actin homolog MreB. Proceedings of the National Academy of Sciences 111, 3585-3590 (2014).

20 Colavin, A., Shi, H. & Huang, K. C. RodZ modulates geometric localization of the bacterial actin MreB to regulate cell shape. Nature communications 9, 1280 (2018).

21 Salje, J., van den Ent, F., de Boer, P. & Löwe, J. Direct membrane binding by bacterial actin MreB. Mol Cell 43, 478-487, doi:10.1016/j.molcel.2011.07.008 (2011).

22 Gitai, Z., Dye, N. A., Reisenauer, A., Wachi, M. & Shapiro, L. MreB actinmediated segregation of a specific region of a bacterial chromosome. Cell 120, 329-341 (2005).

23 Awuni, Y., Jiang, S., Robinson, R. C. & Mu, Y. Exploring the A22-Bacterial Actin MreB Interaction through Molecular Dynamics Simulations. The Journal of Physical Chemistry B 120, 9867-9874 (2016).

24 Shiomi, D. et al. Mutations in cell elongation genes mreB, mrdA and mrdB suppress the shape defect of RodZ-deficient cells. Molecular microbiology 87, 1029-1044 (2013).

25 Van Den Ent, F., Johnson, C. M., Persons, L., De Boer, P. & Löwe, J. Bacterial actin MreB assembles in complex with cell shape protein RodZ. The EMBO journal 29, 1081-1090 (2010).

26 Bendezù, F. O., Hale, C. A., Bernhardt, T. G. & De Boer, P. A. RodZ (YfgA) is required for proper assembly of the MreB actin cytoskeleton and cell shape in E. coli. The EMBO journal 28, 193-204 (2009).

27 Schmidt, A. et al. The quantitative and condition-dependent Escherichia coli proteome. Nature biotechnology 34, 104 (2016).

28 Quint, D. A., Gopinathan, A. & Grason, G. M. Shape selection of surface-bound helical filaments: biopolymers on curved membranes. Biophysical journal 111, 1575-1585 (2016).

29 Deng, Y., Sun, M. & Shaevitz, J. W. Direct measurement of cell wall stress stiffening and turgor pressure in live bacterial cells. Physical Review Letters 107, 158101 (2011).

30 Hussain, S. et al. MreB filaments align along greatest principal membrane curvature to orient cell wall synthesis. eLife 7, e32471 (2018).

31 Ouzounov, N. et al. MreB orientation correlates with cell diameter in Escherichia coli. Biophysical journal 111, 1035-1043 (2016).

32 Hagen, N., Gao, L. & Tkaczyk, T. S. Quantitative sectioning and noise analysis for structured illumination microscopy. Optics express 20, 403-413 (2012).

33 Levin, M. Left-right asymmetry in embryonic development: a comprehensive review. Mechanisms of development 122, 3-25 (2005).

34 Schilthuizen, M. & Davison, A. The convoluted evolution of snail chirality. Naturwissenschaften 92, 504-515 (2005).

35 Wang, J.-S. et al. Hierarchical chirality transfer in the growth of Towel Gourd tendrils. Scientific reports 3, 3102 (2013).

36 Jauffred, L., Vejborg, R. M., Korolev, K. S., Brown, S. & Oddershede, L. B. Chirality in microbial biofilms is mediated by close interactions between the cell surface and the substratum. The ISME journal 11, 1688 (2017).

37 Leaver-Fay, A. et al. in Methods in enzymology Vol. 487 545-574 (Elsevier, 2011).

38 Wang, S. & Wingreen, N. S. Cell shape can mediate the spatial organization of the bacterial cytoskeleton. Biophysical journal 104, 541-552 (2013).

39 Polka, J. K., Kollman, J. M., Agard, D. A. & Mullins, R. D. The structure and assembly dynamics of plasmid actin AlfA imply a novel mechanism of DNA segregation. Journal of bacteriology 191, 6219-6230 (2009).

40 Wan, L. Q. et al. Micropatterned mammalian cells exhibit phenotype-specific left-right asymmetry. Proceedings of the National Academy of Sciences 108, 1229512300 (2011).

41 Pereira, A. R. et al. FtsZ-dependent elongation of a coccoid bacterium. MBio 7, e00908-00916 (2016).

42 Phillips, J. C. et al. Scalable molecular dynamics with NAMD. J. Comput. Chem. 26, 1781-1802, doi:10.1002/jcc.20289 (2005).

43 Best, R. B. et al. Optimization of the additive CHARMM all-atom protein force field targeting improved sampling of the backbone phi, psi and side-chain chi(1) and chi(2) dihedral angles. J Chem Theory Comput 8, 3257-3273, doi:10.1021/ct300400x (2012).

44 Mackerell, A. D., Jr., Feig, M. & Brooks, C. L., 3rd. Extending the treatment of backbone energetics in protein force fields: limitations of gas-phase quantum mechanics in reproducing protein conformational distributions in molecular dynamics simulations. J. Comput. Chem. 25, 1400-1415, doi:10.1002/jcc.20065 (2004).

45 Jorgensen, J. H., Johnson, J. E., Alexander, G. A., Paxson, R. & Alderson, G. L. Comparison of automated and rapid manual methods for the same-day identification of Enterobacteriaceae. Am. J. Clin. Pathol. 79, 683-687 (1983).

46 Tuckerman, M., Berne, B. J. & Martyna, G. J. Reversible multiple time scale molecular dynamics. J. Chem. Phys. 97, 1990-2001 (1992).

47 Brünger, A., Brooks, C. L. & Karplus, M. Stochastic boundary conditions for molecular dynamics simulations of ST2 water. Chem Phys Lett 105, 495-500 (1984).

48 Feller, S. E., Zhang, Y., Pastor, R. W. & Brooks, B. R. Constant pressure molecular dynamics simulation: the Langevin piston method. J. Chem. Phys. 103, 4613-4621 (1995).

49 Humphrey, W., Dalke, A. & Schulten, K. VMD: visual molecular dynamics. J. Mol. Graph. 14, 33-38, 27–28 (1996).

50 Hsin, J., Gopinathan, A. & Huang, K. C. Nucleotide-dependent conformations of FtsZ dimers and force generation observed through molecular dynamics simulations. Proc Natl Acad Sci U S A 109, 9432-9437, doi:10.1073/pnas.1120761109 (2012).

51 Pettersen, E. F. et al. UCSF Chimera—a visualization system for exploratory research and analysis. Journal of computational chemistry 25, 1605-1612 (2004).

52 Zoete, V., Cuendet, M. A., Grosdidier, A. & Michielin, O. SwissParam: a fast force field generation tool for small organic molecules. Journal of computational chemistry 32, 2359-2368 (2011).

53 Webb, B. & Sali, A. Comparative protein structure modeling using MODELLER. Current protocols in protein science 86, 2.9. 1–2.9. 37 (2016).

54 Ursell, T. et al. Rapid, precise quantification of bacterial cellular dimensions across a genomic-scale knockout library. BMC biology 15, 17 (2017).

55 Sliusarenko, O., Heinritz, J., Emonet, T. & Jacobs-Wagner, C. High-throughput, subpixel precision analysis of bacterial morphogenesis and intracellular spatiotemporal dynamics. Molecular microbiology 80, 612-627 (2011).

